# Data-driven reduction of dendritic morphologies with preserved dendro-somatic responses

**DOI:** 10.1101/2020.04.06.028183

**Authors:** Willem A.M. Wybo, Jakob Jordan, Benjamin Ellenberger, Ulisses M. Mengual, Thomas Nevian, Walter Senn

## Abstract

Dendrites shape information flow in neurons. Yet, there is little consensus on the level of spatial complexity at which they operate. We present a flexible and fast method to obtain simplified neuron models at any level of complexity. Through carefully chosen parameter fits, solvable in the least squares sense, we obtain optimal reduced compartmental models. We show that (back-propagating) action potentials, calcium-spikes and NMDA-spikes can all be reproduced with few compartments. We also investigate whether afferent spatial connectivity motifs admit simplification by ablating targeted branches and grouping the affected synapses onto the next proximal dendrite. We find that voltage in the remaining branches is reproduced if temporal conductance fluctuations stay below a limit that depends on the average difference in input impedance between the ablated branches and the next proximal dendrite. Further, our methodology fits reduced models directly from experimental data, without requiring morphological reconstructions. We provide a software toolbox that automatizes the simplification, eliminating a common hurdle towards including dendritic computations in network models.

## Introduction

Morphological neuron models have been instrumental in neuroscience^1^. Major experimental discoveries, for instance that N-methyl-D-aspartate (NMDA) channels^2^ produce local dendritic all or none responses^3,4^, or that dendritic Ca^2+^-spikes mediate coincidence detection between distal inputs and somatic action potentials (APs)^5^, have been combined in morphological models to arrive at a consistent picture of dendritic integration: the dendrite is an intricate system of semi-independent subunits^6,7^, amenable to dynamic regulation ^8,9^, and able to distinguish specific input patterns^10,11^.

Nevertheless, morphological models are not without shortcomings. They are highly complex and consist of thousands of coupled compartments, each receiving multiple non-linear currents. The parameters of the models, typically fitted with evolutionary algorithms to electro-physiological recordings^12–14^, number in the tens of thousands. Since recordings can only be obtained at a few dendritic sites, these fits are under-constrained, and thus susceptible to over-fitting. Accordingly, the single-neuron fitting challenge, where model performance was measured on unseen spike trains, was not won by a biophysical model, but by an abstract spiking model^15^. Finally, many network-level observations can be explained without morphological models^16^.

These shortcomings underline the need to find the essential computational repertoire of a neuron: the set of computations needed to understand brain functions such as learning and memory. Model simplification is crucial in this endeavour, as it elucidates the lowest level of complexity at which computational features are preserved. Conceptually, the simplification thus extracts the essential elements required for the computation from the underlying biophysics. Simulation-wise, the reduced model requires few resources, and can thus be integrated in large-scale networks. Experimentally, the reduced model likely leads to a well-constrained fit.

Past simplification efforts can be grouped in two categories: approaches that use traditional compartments, but with adapted parameters^17–22^, and approaches that rely on more advanced mathematical techniques^23,24^. Many authors apply ad-hoc morphological simplifications and then adjust model parameters to conserve geometrical quantities, such as surface area^21,25,26^, or electrical quantities, such as attenuation^18^ and transfer impedance^22^. Other authors propose two-compartment models whose parameters are adjusted to reproduce certain response properties^17,27^. Nevertheless, these approaches often lack the flexibility to be adapted to a wide range of dendritic computations. Further, particular afferent spatial connectivity motifs might be lost, so that essential computations are not captured by such a reduction. Hence, in a network model, the computational relevance of these motifs might unknowingly be ignored. Advanced mathematical techniques on the other hand are very flexible, as they can incorporate the response properties of the morphology implicitly^23,24^. However, these techniques are not supported by standard simulation software, such as NEURON^28^, a considerable hurdle towards their integration in the canonical neuroscience tool-set.

Here, we introduce a simplification method with the flexibility of advanced mathematical techniques, but using traditional compartments. The approach accommodates any morphological model in public repositories^29^ and commonly used ion channels^30^. The reduced models represent the optimal approximation to the dendritic impedance matrix, evaluated at any spatial resolution of choice. We fit ion channels by approximating the quasi-active impedance matrices^31^. All our fits are uniquely solvable linear least-squares problems. The obtained models extrapolate well to non-linear dynamics, and reproduce back-propagating APs (bAPs), Ca^2+^-spikes^5^ and NMDA spikes^3,4^ with few compartments. Additionally, we investigate whether a dendritic tree with given afferent spatial connectivity motifs can be simplified by ablating subtrees or branches and grouping synapses in the next proximal compartment. We find that effective weight-rescale factors for synapses can be computed if temporal conductance fluctuations stay below a limit that depends the difference in input impedance between the ablated branch and the next proximal dendrite. Under application of these factors, voltage responses in the simplified tree are preserved. Finally, we demonstrate that our approach can fit reduced models to experimental recordings, without the need to reconstruct full morphologies. We have created a Python toolbox (NEural Analysis Toolbox – NEAT – https://github.com/unibe-cns/NEAT) that implements this method together with an extension to NEURON to simulate the obtained models.

## Results

### A systematic simplification of complex neuron morphologies

Our simplification strategy fits compartments with active channels to a reduced set of dendritic locations of interest (Fig 1A, left-middle). The locations, together with the morphology, provide a tree structure for the reduced model (Fig 1A, middle – right) where each node corresponds to a traditional compartment and each edge to a coupling conductance. The fit requires that the reduced model responds similarly to perturbative current steps as the full neuron, at the set of chosen locations (Fig 1B). We consider perturbations around any spatially uniform holding potential *v_h_*. Experimentally, *v_h_* may be reached by injecting a constant current *i_h_*, and thus *v_h_* is the effective equilibrium potential under this current injection. In models, we do not need to know *i_h_*; we simply assume that *v_h_* is the equilibrium potential around which to linearize the model. Note that if *i_h_* = 0, *v_h_* is the resting membrane potential.

**Figure 1.**
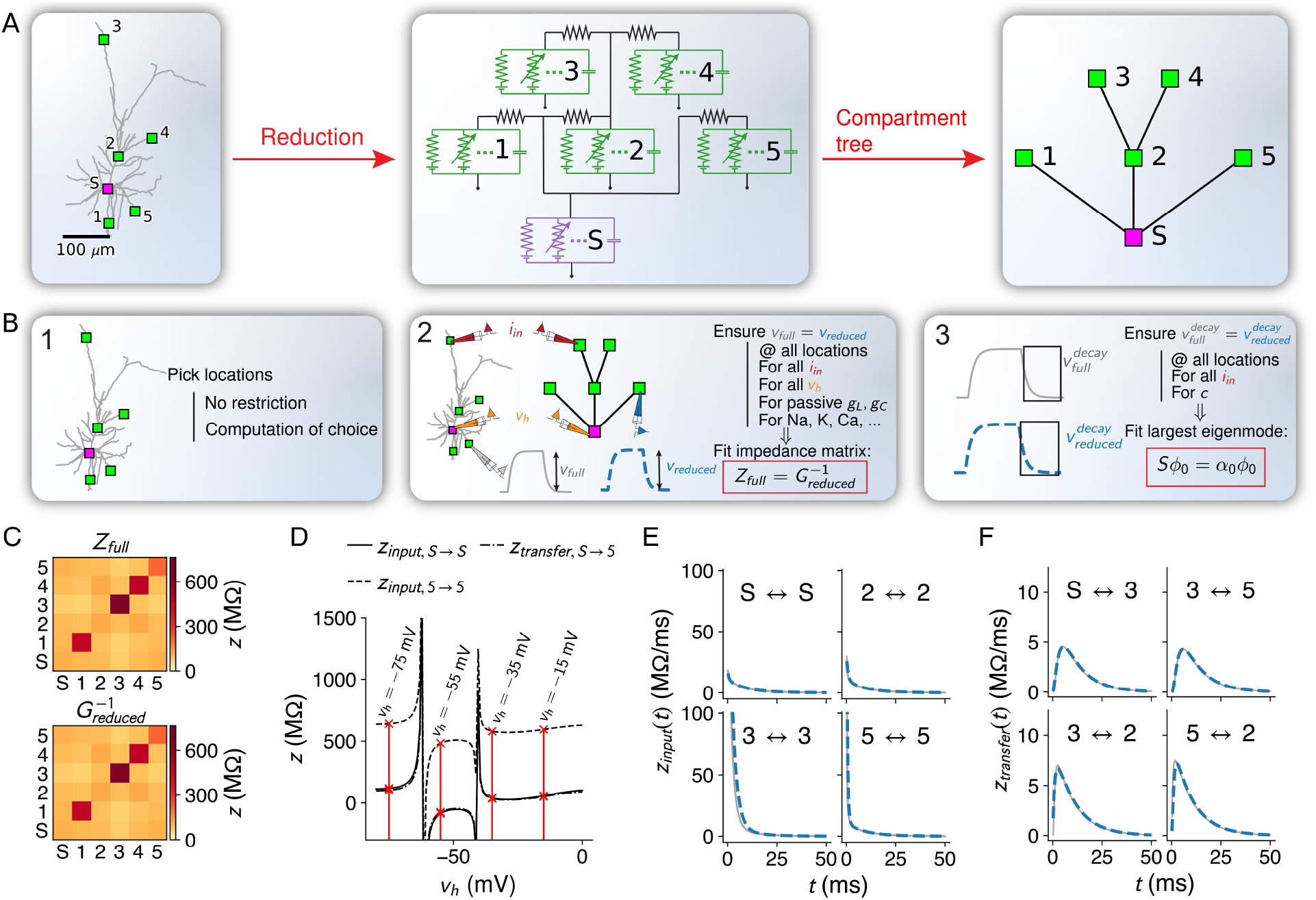
Flexible and accurate reduction methodology. **A:** For any set of locations on a given morphology (left, here an L2/3 pyramidal cell^10^), a reduced compartmental model can be derived (middle), with an associated schematic representation (right). **B:** Steps of our approach: (1) choice of locations at which the reduced model should reproduce the full model’s voltage, (2) coupling, leak and channel conductances are fitted to impedance matrices derived from the full model at different holding potentials and (3) capacitances are fitted to mimic the largest eigenmode of the full model. **C:** The full model’s impedance matrix (top) restricted to the 5 locations (A) is approximated accurately by the inverse of the conductance matrix of the reduced model (bottom). Labels correspond to locations in A. **D:** Example components of the quasi-active impedance matrix of the full model, equipped with a Na-channel, as a function of the holding potential *v_h_*. Red lines show the four holding potentials at which our methodology evaluates the impedance matrix. Singularities correspond to holding potentials where the linearization is invalid, and should be avoided in the fit. Labels correspond to locations in A. **E:** Temporal shape of exemplar input impedance kernels of the full model (grey) and their reduced counterparts (blue, dashed). **F:** Same as in E, but for transfer impedance kernels.

Suppose we have chosen *N* locations. Let *δ*_**i**_ be a vector describing perturbative input currents to each of those *N* locations. The linearized voltage response of the full neuron, at those locations, is

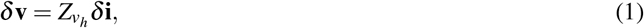

with *Z_v_h__* the quasi-active^31^ *N × N* impedance matrix. In experiments, *Z_v_h__* is extracted from *δ*_**v**_ measured in response to *δ*_**i**_. In models, *Z_v_h__* is algorithmically computed based on the full morphology and the dynamics of the membrane conductances^31^. The *N × N* conductance matrix *G_v_h__* for the reduced model contains on its diagonal the sum of the unknown leak, coupling and linearized ion channel conductance parameters for each compartment, and in its non-zero off-diagonal entries the unknown coupling conductance parameters between connected compartments. This matrix relates the reduced model’s voltage response to the perturbative input

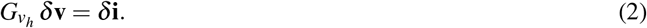

The full neuron and reduced model will behave similarly for all possible perturbative inputs *δ*_**i**_ if their responses *δ*_**v**_ match. From (1) and (2), it follows that *Z_v_h__* should be the inverse of *G_v_h__*. Consequently, we require that multiplying the known *Z_v_h__* (measured or calculated) by the parametric *G_v_h__* yields the identity

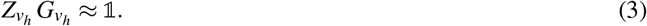

As this system of equations is linear in leak, coupling and channel conductances of the reduced compartments, it can be cast into a least squares problem and solved accurately (Fig 1C, Fig S1A).

Since ion channel activation depends on voltage, *Z_v_h__* changes with *v_h_* (Fig 1D). The fit must disentangle the changes in *Z_v_h__* induced by the various channels. In models, we first block all ion channels in the full and reduced models and fit Gpas to Zpas according to (3), and thus obtain leak and coupling conductances for each compartment. Then, we unblock one ion channel at a time and decompose the conductance matrix into *G_v_h__* = *G*_pas_ + *G*_*v*_*h*_,chan_, with *G*_*v*_*h*_,chan_ a diagonal matrix containing the conductances of the unblocked channel, linearized around *v_h_*, at each compartment. Thus *G*_*v*_*h*_,chan_ depends on the unknown maximal ion channel conductance parameters. With *Z_v_h__* and *G*_pas_ known, we optimize these maximal conductances so that the left hand side of

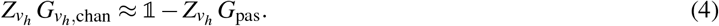

matches the right hand side, and that it does so for multiple holding potentials (we chose *v_h_* = −75, −55, −35 and −15 mV, Fig 1D). Thus, we obtain an overdetermined system of equations in the unknown maximal channel conductances at each compartment, and compute its solution in the least mean squares sense.

To fit capacitances for each compartment it suffices to consider passive membrane dynamics. Blocking all ion channels in the reduced model (*G*_*v*_*h*_,chan_ = 0), temporal voltage responses follow

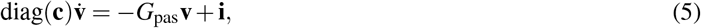

with **c** containing the unknown capacitances of each compartment. We require that voltage-decay back to rest, as described by (5), matches voltage-decay in the full neuron with all active channels blocked, at the *N* chosen locations. According to linear dynamical systems theory^32^, this decay can be decomposed as a sum of exponentially decaying eigenmodes, each with an associated eigenvalue *α_k_* and eigenvector ***ϕ***_*k*_. The former gives the exponential time-scale *τ_k_* = 1/*α_k_* of the decay and the latter the spatial profile of the mode. The most prominent of these modes has the largest time-scale *τ*_0_ = 1/*α*_0_, and primarily models loss of voltage through the membrane^33^. In experiments, we extract this mode by fitting an exponential to the voltage decay. In full models, we compute the eigenvalue *α*_0_ and eigenvector ***ϕ***_0_ – restricted to the *N* chosen locations – with the separation-of-variables method^34^. In the reduced model, eigenmodes are found as the eigenvalues and -vectors of the matrix *S* = -diag(**c**)^-1^ *G*_pas_. To fit the capacitances, we require that *S* has *α*_0_ as eigenvalue and ***φ***_0_ as corresponding eigenvector:

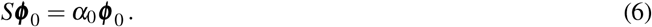

This system of equations is linear in the reciprocals of the capacitances entering in S and can thus be solved efficiently. To accurately reproduce the spatio-temporal voltage responses of the full model, the reduced model must approximate the “impedance kernels^24^” *z*(*t*), the inverse Fourrier transforms of the frequency dependent elements of the impedance matrix. Despite only fitting the largest time-scale 1/*α*_0_, we accurately reproduce kernels at all times (Fig 1E-F, Fig S1B).

Finally, we fit leak reversals for each compartment so that the resting voltage matches the full models resting voltage, at the *N* chosen locations. To do so, we evaluate all ion channel currents in the reduced model at the full model’s resting voltage and require that temporal voltage derivatives are zero. The obtained equations are linear in the reduced model’s leak reversals, and can be solved efficiently.

### Reduced models match the voltage response of their full counterparts

We demonstrated the reduction on two computations that require accurate spatio-temporal interaction. We reproduced sequence discrimination in an L2/3 pyramidal cell model^10^, where a neuron responds more strongly to a centripetal sequence of inputs than to a centrifugal one, by only retaining compartments on a single branch (Fig 2A). We also reproduced input-order detection^35^, where a neuron responds more strongly when one input arrives before the other, by only retaining compartments at the soma and the two input sites (Fig 2B).

**Figure 2.**
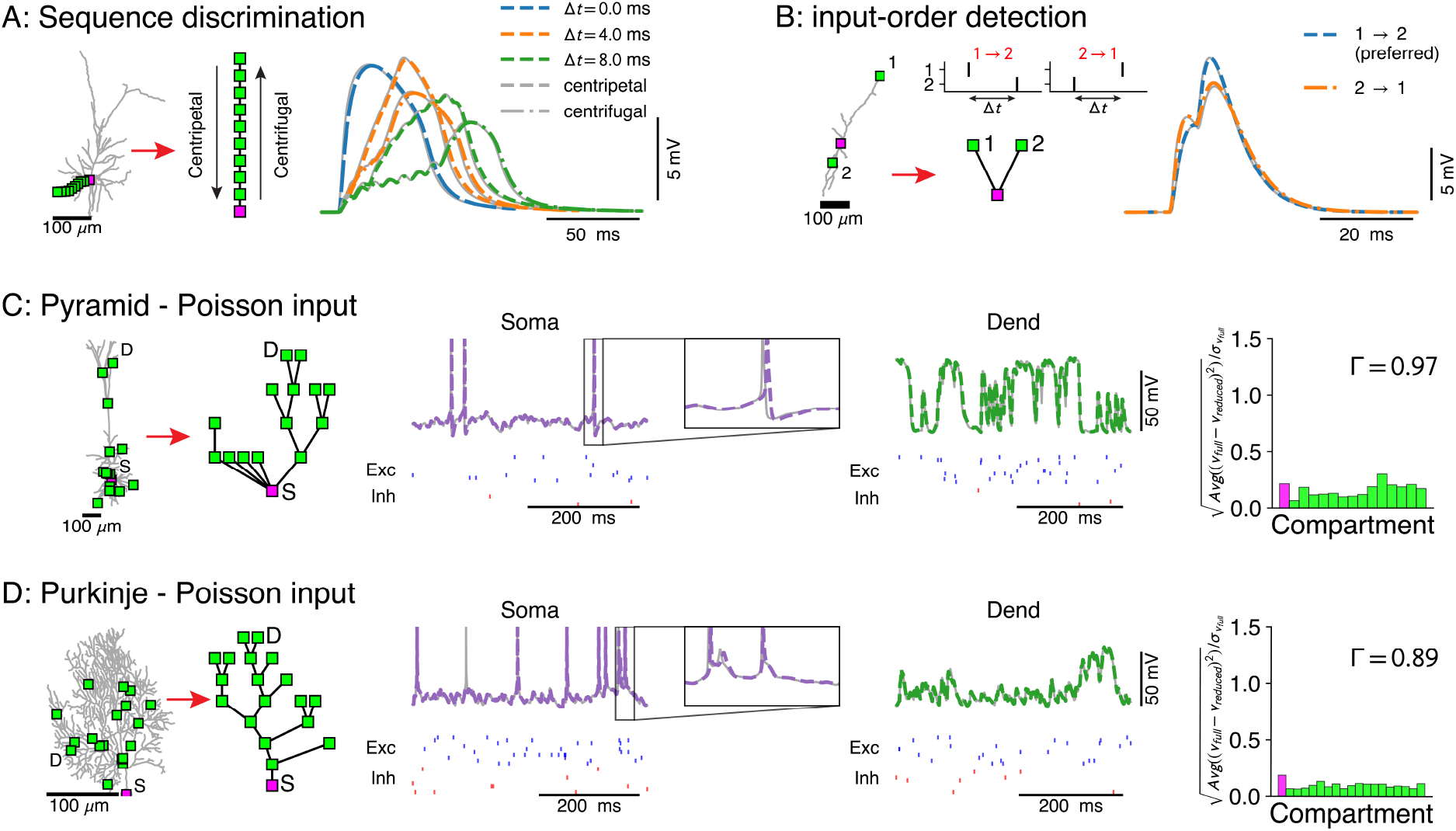
Voltage-match between full and reduced models for spatio-temporal dendritic computations. **A:** Reduction of full model (right) to a single branch (middle) reproduces sequence discrimination (right, full model in grey and reduced model colored for different time-steps between inputs, centripetal (dashed) and centrifugal (dash-dotted)). **B:** Full model (left) and three-compartment reduction (middle, bottom) discriminate temporal order of inputs, where the response to inputs (middle, top) ordered 1 → 2 is stronger than 2 → 1. Voltage responses (right) in full model (grey) and reduced model (colored). **C, D:** Reductions of resp. L5 pyramidal and Purkinje cells with active ion channels at the soma, and excitatory (AMPA+NMDA) and inhibitory (GABA) synapses at dendritic compartment sites. From left to right: full model with compartment sites (soma *S* and a selected dendrite site *D* are labeled), reduced model, somatic voltage with zoom on a single AP (full model in grey and reduced model in purple, input spikes at the bottom), dendritic voltage at D (full model in grey and reduced model in green, input spikes at the bottom) and the average root mean square voltage errors, normalized by voltage standard deviation, at each compartment. Spike coincidence factor **Γ** is also shown.

We then tested our approach with non-linear currents concentrated at discrete points along the morphology. In two biophysical models – an L5 neocortical pyramid^12^ (Fig 2C) and a Purkinje cell^36^ (Fig 2D) – we removed active dendritic channels, but added their total conductance at rest to the passive leak (we term this the “passified” model), while retaining all active channels at the soma. The soma, with its AP-generating channels, is such a point. We clustered sets of excitatory (*α*-amino-3-hydroxy-5-methyl-4-isoxazolepropionic acid (AMPA) + NMDA) and inhibitory (*γ*-aminobutyric acid (GABA)) synapses at randomly selected points. For both models, membrane voltage traces were reproduced, both at the soma and at dendritic sites (middle + right panels), and at least 89% of APs coincided within a 6 ms window^37^.

What level of spatial complexity reproduces characteristic responses generated by somatic and dendritic ion channels? These responses arise through the interplay of integrative properties of the morphology – captured in the impedance kernels *z*(*t*) – and channel dynamics. If a full model (here the L2/3 pyramid) is reduced to a single somatic compartment (Fig 3A), the impedance kernel has only one time-scale – the membrane time-scale (Fig 3B, top – purple). The kernel of the full model contains additional shorter time-scales (Fig 3B, top – grey), leading to a faster AP-onset than in the reduced model (Fig 3B, bottom). This effect is thus a fundamental limitation of the single-compartment reduction. Extending the compartmental model with 4 nearby dendritic sites (Fig 3A) adds fast components to the somatic impedance kernel (Fig 3B, top – green) that model the spread of voltage to the other compartments. These components increase AP amplitude and decrease AP delay (Fig 3C), leading to a better match with the full model.

**Figure 3:**
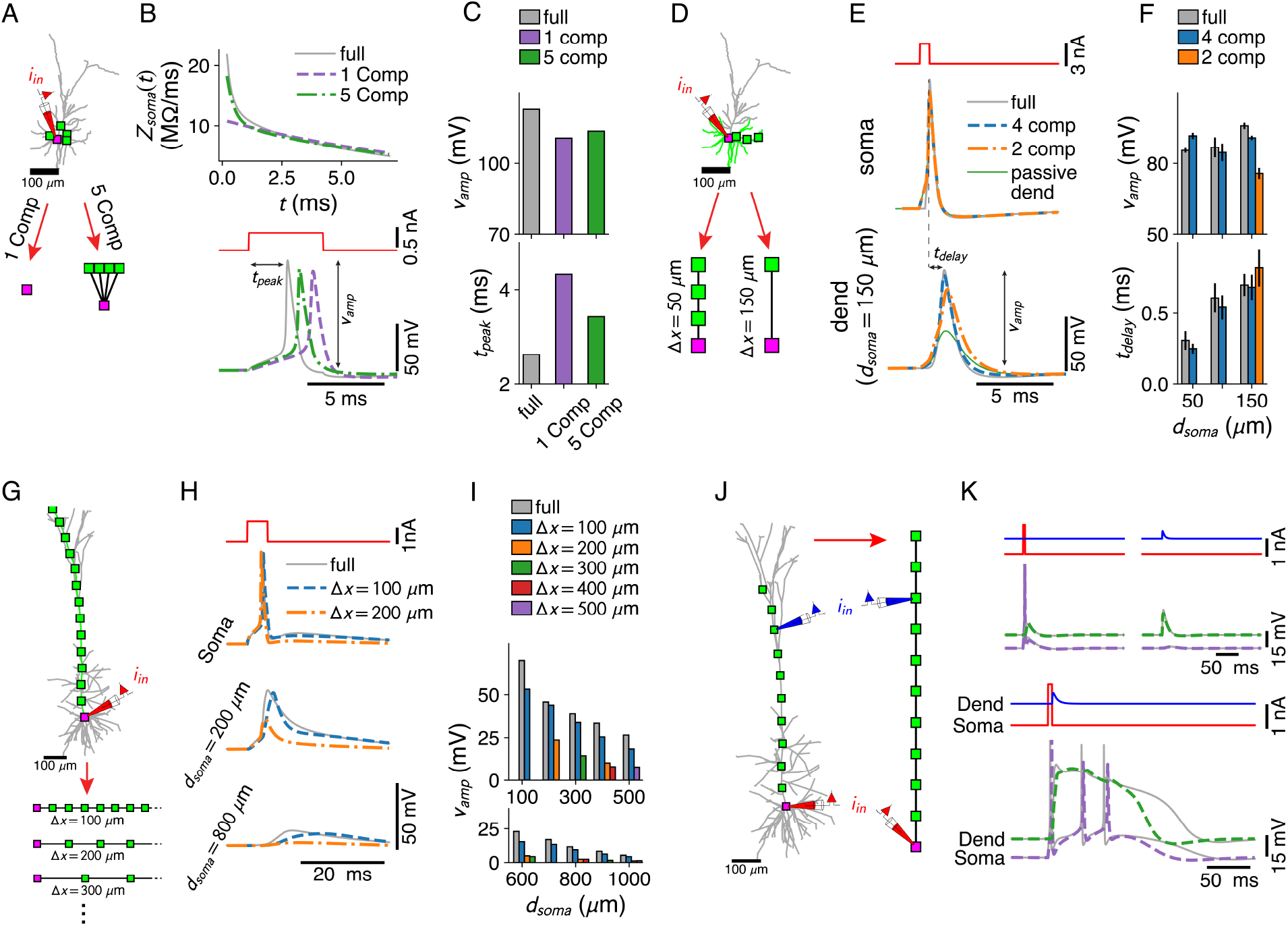
Dendritic computations with active channels are captured by our reduced models. **A-C:** Effect of compartment distribution on AP dynamics in reduced models. **A:** One- and five-compartment reductions of the L2/3 pyramid, equipped with somatic and dendritic ion channels. **B:** Differences in short-timescale behaviour in somatic input impedance kernels (top) between full model (grey), and one- (purple) and five-compartment (green) reductions result in different AP delays (bottom). **C:** AP amplitude (top) and AP delay (bottom) for the three models (colors as in B). **D-F:** Effect of compartment distribution on AP backpropagation in basal dendrites. **D:** Four- and two-compartment reductions of a basal branch. **E:** APs at soma (top) and most distal compartment site (bottom) for four models (full in grey, four compartments in blue, two compartments in orange, and full but with a passive dendrite in green). **F:** Amplitude (top) and delay (bottom) for bAPs at different distances from soma (if compartment is present in model), averaged over all basal branches longer than 150 *μ*m (error bars indicate standard deviation). **G-I:** AP backpropagation in the apical dendrite of the L5 pyramid. **G:** Reductions of the apical dendrite with increasing inter-compartment spacing. **H:** Voltage waveform at soma (top, full in grey, Δ*x* = 100 μ*m* in blue, Δ*x* = 200 *μm* in orange) and two dendritic sites (middle, bottom). **I**: Waveform amplitude as a function of distance to soma for various inter-compartment spacings. **J, K:** Ca^2+^ -spike mediated coincidence detection. **J:** Reduction of the L5-pyramid’s apical dendrite to 11 compartments. **K**: Response to a somatic current pulse (top, left), a dendritic synaptic current waveform (top, right) and the coincident arrival of both inputs (bottom).

Dendritic ion channels support the back-propagation of APs^38^. For each basal branch of at least 150 μm in the L2/3 pyramid, we derived reduced models with three compartments (at 50, 100 and 150 μm from the soma) and with one compartment. We compared bAP amplitudes and delays relative to the somatic AP (Fig 3E) and found that even models with a single distal compartment support active back-propagation (Fig 3E, F). In the apical dendrite of the L5 pyramid (Fig 3G), we found that a distance step of 100 μm between compartments was required to support bAPs to the same degree as in the full model (Fig 3H, I).

Finally, we considered the pairing of a somatic current pulse with a dendritic post-synaptic potential (PSP) waveform. We found that an 11-compartment reduction (Fig 3J) – with compartments spaced at 100 *μ*m through the apical trunk to support bAPs until the Ca^2+^ hot-zone – reproduced the Ca^2+^-mediated coincidence detection mechanism^5^.

### Conditions under which afferent spatial connectivity motifs can be simplified

Our method allows us to reduce morphological complexity by removing any branch without affecting the integrative properties of other branches. We consider the computational repertoire of the neuron intact under such an ablation if we can shift the synapses on the ablated branch to the nearest intact proximal compartment, rescale their weights with a temporally constant factor *β* and recover the original voltage at intact sites (Fig 4A). Which spatial synapse distributions admit simplification in this sense, and under which input conditions? For a single, current-based synapse, we analytically compute the weight-rescale factor *β*_curr_ = *z_cs_*/*z_cc_*, with *z_cs_* the transfer impedance from synapse site *s* to compartment site *c* and *z_cc_* the input impedance at *c*. Since *z_cs_* < *z_cc_*^39^, the weight-rescale factor weakens the shifted synapse so that it matches the attenuation of its distal counterpart. Nevertheless, *β*_curr_ is often close to one, and hence negligible (Fig 4B, C). For a conductance-based synapse, rescaling weights by *β*_curr_ does not accurately fit the full model (Fig 4B). Decomposing the synaptic conductance *g*(*t*) as *g* + *δg*(*t*), with *g* the temporal average and *δg*(*t*) the fluctuations around *g*, we analytically compute that the weight-rescale factor for a conductance-based synapse

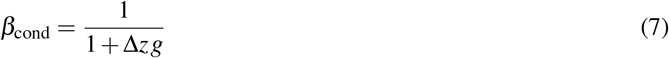

recovers the full model’s voltage if

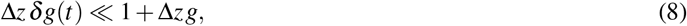

where Δ*z* = *z_ss_* – *z_cc_* is the difference between input impedance *z_ss_* at the original synapse site *s* and input impedance *z_cc_* at the nearest proximal compartment site *c* (see methods). Using *β*_cond_ weakens the synapse so that voltage at all compartments more proximal than *c* and in their side branches (the “downstream voltage”) is reproduced (Fig 4D).

**Figure 4.**
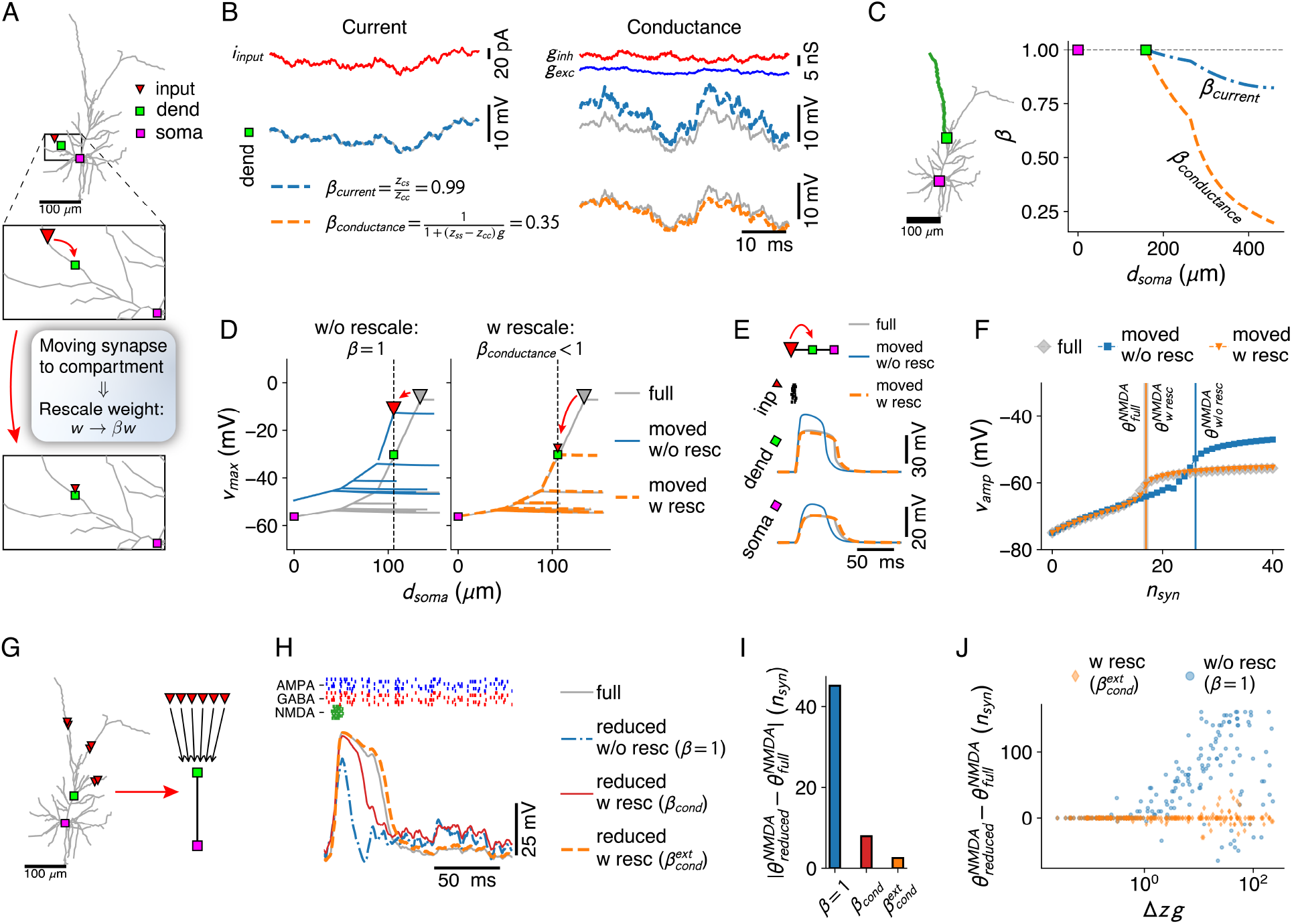
Simplification of afferent spatial connectivity motifs. **A:** Removal of a branch with a synapse (red triangle) is considered possible if the correct voltages at the compartment sites (here, dendritic – green square and somatic – purple square) can be obtained by shifting the synapse to the compartment site and rescaling its weight with a fixed factor *β* for the all input conditions under consideration. **B:** Comparison between weight-rescale factors for current-based (left) and conductance-based (right) input. Voltage trace for dendritic compartment without rescaling (grey) and with the current-based scale factor *β*_curr_ compensating for attenuation (blue). Bottom right panel shows voltage trace with conductance-based scale factor *β*_cond_. **C:** Current- and conductance based scale factors in the green dendritic branch, for a shift of a synapse at a given distance from the soma to the dendritic compartment site (green square). **D:** Spatial peak voltage without (left, blue) and with (right, orange) application of *β*_cond_. **E:** For a cluster of AMPA+NMDA synapses in isolation, scale factors for physiological constants can be obtained (see methods) that reproduce the correct voltage waveform. Colors as in D. **F:** Maximal amplitude of NMDA spike waveform upon activation of increasing number of synapses – NMDA spike thresholds indicated with vertical lines. Colors as in D. **G-H:** Removing a whole subtree and shifting multiple synapses (red triangles) to the next proximal compartment site (green square). NMDA spike generation (grey voltage trace) at the compartment site through burst inputs to local AMPA+NMDA synapses (green inputs), with AMPA (blue) and GABA (red) background inputs spread throughout the subtree. Reductions shown without rescaling (blue, dash-dotted), with the analytical single-site rescaling rule (red, full) and the numerical multi-site rule (orange, dashed). **I:** Error in NMDA spike threshold for the three cases in **H. J:** Dependence of the error in NMDA-spike threshold on the factor Δ*zg*, with Δ*z* the average input impedance difference between synapse sites and the compartment and g the average synaptic conductance.

If *s* and *c* are close, Δ*z* is small and (8) is satisfied for plausible conductance fluctuations. Thus, shifting the synapse to the next proximal compartment keeps the voltage response intact. If the separation between *s* and *c* increases, ***Δ****z* increases proportionally, and the magnitude of tolerated conductance fluctuations shrinks (8). Thus, for a given magnitude of ***δ**g*(*t*), it is possible that a thick branch can be removed, but not a thin branch, as the input impedance increase is less steep in the former. Nevertheless, (8) suggests that even for large ***Δ****z* the full model’s voltage can be recovered by downscaling synaptic weight by ***β***_cond_, but only if *δg*(*t*) ≪ *g*. A tonic level of AMPA and/or GABA activation, as in the high-conductance state^40^, can implement this input condition. For NMDA synapses, this input condition does not hold, as NMDA spikes arise through a strong, transient conductance increase. However, when a cluster of such synapses – to be moved to a proximal site – is considered in isolation, a workaround can be found by applying constant rescale factors to threshold and width of the Mg-block, as well as to the reversal (see Methods). These factors recover NMDA spike shape (Fig 4E) and threshold (Fig 4F) when the cluster is moved to from *s* to *c*.

Our analytical weight-rescale factor *β*_cond_ treated reductions where a single site s was moved to *c*. We consider now the ablation of a whole subtree and the shift of all its synapses – 100 AMPA and 100 GABA synapses activated with a Poisson rate – to *c* (Fig 4G). Voltage at *c*, as quantified by the shape (Fig 4H) and threshold (Fig 4I) of NMDA-spikes elicited by inputs at *c*, was not preserved with the analytical *β*_cond_. We numerically extend *β*_cond_ to multi-site inputs (see methods). These extended weight-rescale factors 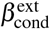 reproduced local voltage (Fig 4H) and threshold (Fig 4I). To elucidate whether 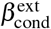 can still be understood as depending on Δ*zg*, we sampled 200 sets of synapses – 100 AMPA and 100 GABA synapses randomly distributed on the subtree – and Poisson activation rates. We found a strong dependence of the error on Δ*zg*, that was corrected by applying 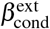 (Fig 2J).

In conclusion, we find that a dendritic branch or subtree can be removed if *δg*(*t*) ≪ *g* by applying the constant weight-rescale factors. Otherwise, the branch can only be removed if Δ*z* is small, so that Δ*z δg* remains below 1 + Δ*zg* for all relevant activation levels.

### Active and passive reduced dendrites under synaptic bombardment

We next investigated how many compartments are needed to reproduce somatic and dendritic responses under a synaptic bombardment, and quantified the contribution of active dendritic Na^+^, Ca^2+^ and K^+^ channels. We mimicked the in-vivo state in the basal dendrites of the L2/3 pyramid by distributing 200 AMPA and 200 GABA synapses receiving Poisson inputs and 20 clusters of AMPA+NMDA synapses receiving bursts of inputs (Fig 5A). We sampled 10 different sets of meta-parameters governing synaptic activation (Poisson and burst rates and synaptic weights, see methods), resulting in output spike rates between 0 and 8 Hz (Fig 5B). We derived reduced models, once based on the full model with all active channels (‘active reduced dendrite’ in Fig 5C) and once based on the passified full model (‘passive reduced dendrite’ in Fig 5C). We distributed increasing numbers of compartment sites on the basal branches and measured somatic and dendritic voltage responses for the active full model and for the reduced models with active and passive dendrites (Fig 5C, see Fig S2 for dendritic traces). Spike coincidence (Fig 5D), subthreshold somatic voltage error (Fig 5E) and dendritic voltage error (Fig 5F) showed a significant improvement when compartment numbers increased from 0 (point neuron – top row in Fig 5C) to 2 per 100 *μ*m. In the presence of active channels the error decreases until up to 3 compartments per 100 *μ*m (Fig 5E, F). Spike coincidence factors were consistently ~10% higher than in the passive case (Fig 5D).

**Figure 5.**
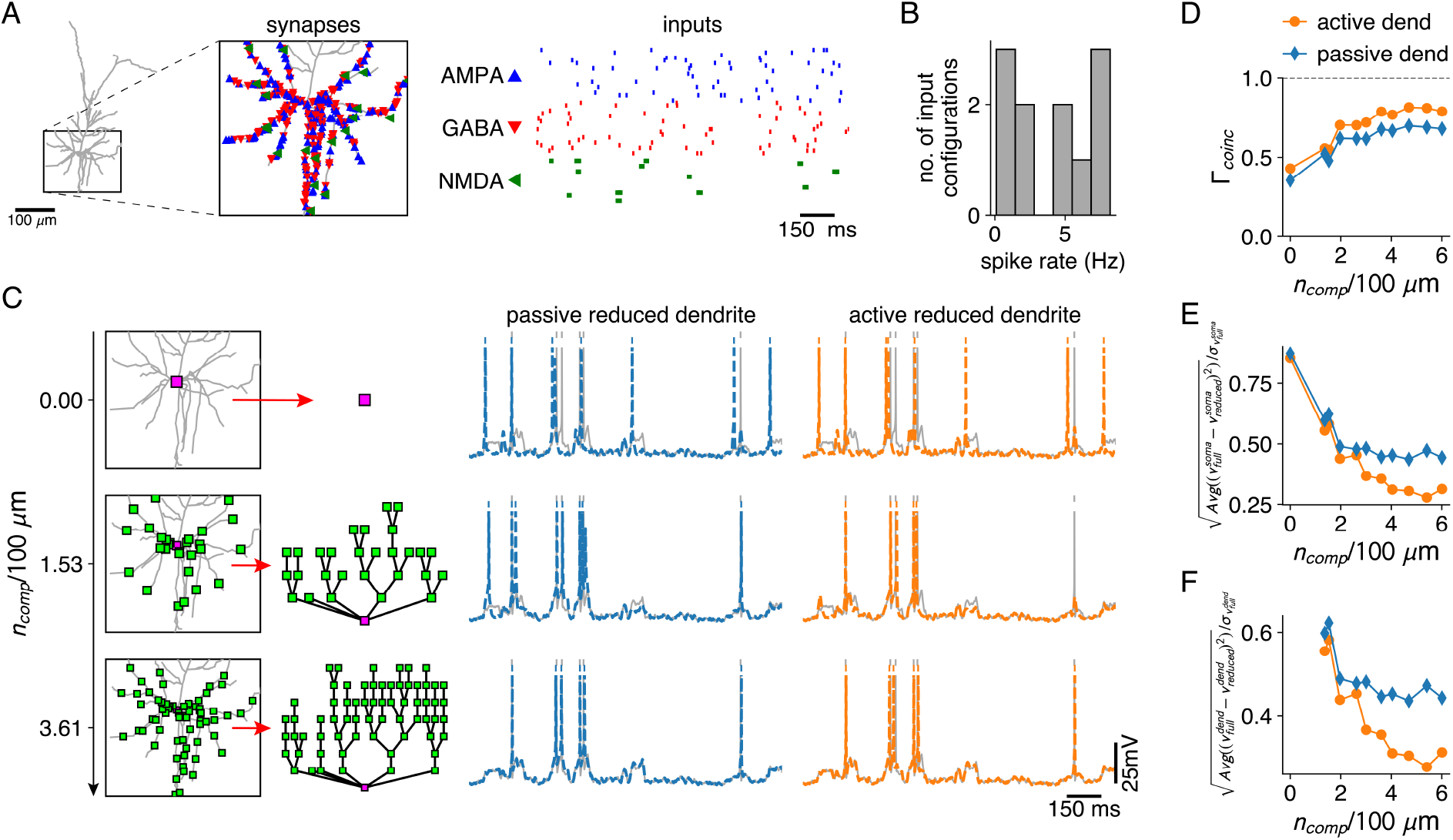
Reductions with active and passive dendritic compartments under in-vivo like conditions. **A:** AMPA and GABA synapses, and AMPA+NMDA synapse clusters are spread randomly throughout the basal dendrites of the L2/3 pyramid (inset). AMPA and GABA synapses receive Poisson inputs and AMPA+NMDA synapses receive bursting input (right). **B:** Output spike rates of the full model for 10 different input configurations. **C:** Reductions with increasing numbers of compartment sites along the basal dendrites (left). Somatic voltage traces and spike times for the full model (grey), for a reduction with passive dendrites (middle, blue) and a reduction with active dendrites (right, orange). **D-F:** Spike coincidence factors (D), relative somatic voltage errors (E) and relative dendritic voltage errors (F) for reductions with increasing numbers of compartments, and with passive (blue) and active dendrites (orange), averaged over all 10 input configurations.

### Fitting reduced models directly to experimental data

Traditionally, creating a morphological neuron model involves a reconstruction of the morphology and an evolutionary fit of the electrical parameters^12–14^. Our method allows skipping these resource-intensive steps (Fig 6A). We adapt our method to a common experimental paradigm where hyper- and depolarizing current steps are injected under applications of various ion channel blockers. We extract impedance matrices and holding potentials from voltage step amplitudes (Fig 6B), and the time-scale of the largest eigenmode from the average voltage decay back to rest (Fig 6C). If the modelling goal is a reduced model, the traditional reconstruction and subsequent reduction both introduce errors (Fig 6D). Thus, we find that while our reduction of the full model is faithful within an expected accuracy margin (Fig 6E), fitting a reduce model directly to the experimental traces is more accurate (Fig 6F).

**Figure 6.**
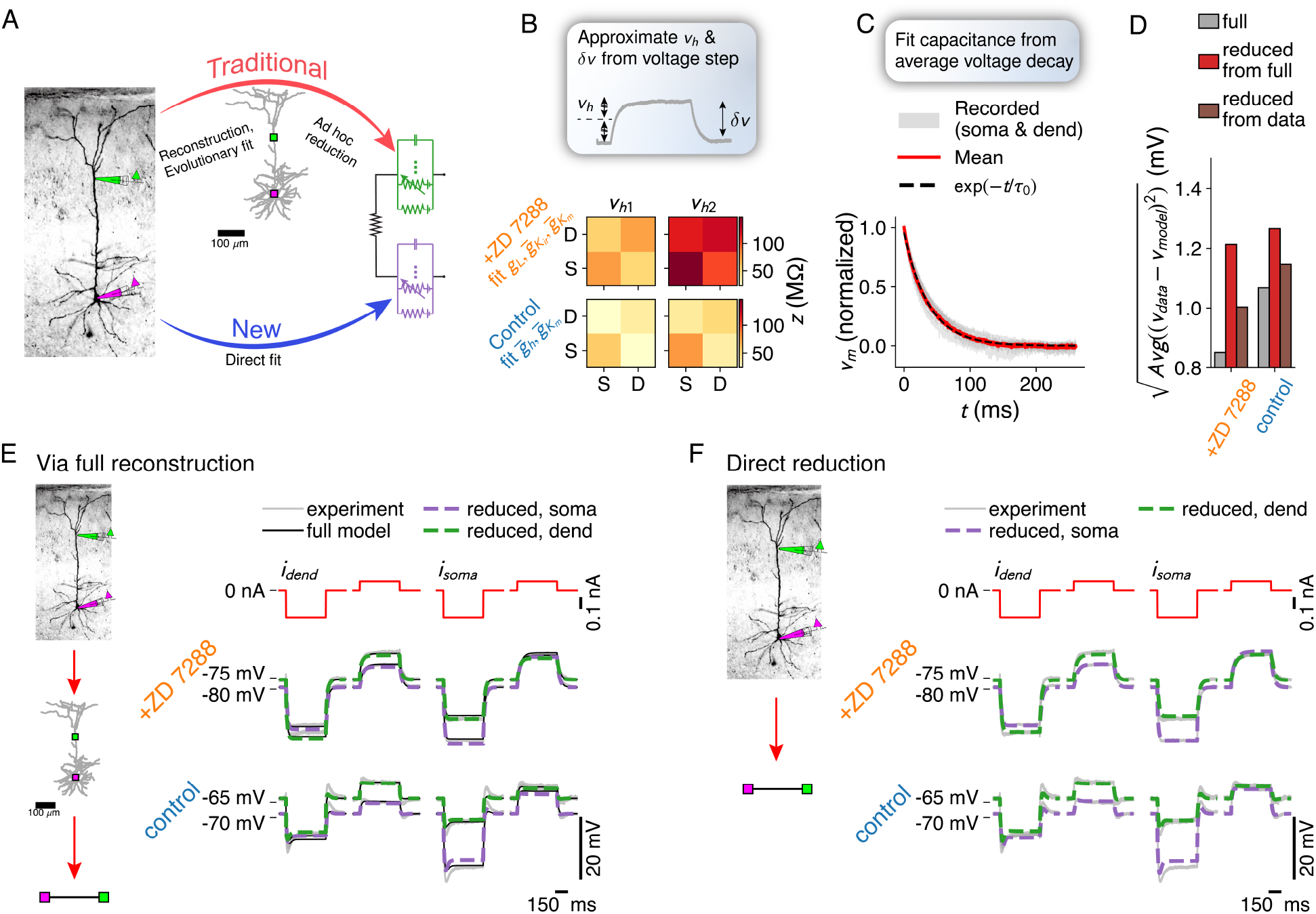
Fitting reduced models directly to experimental data. Comparison between the traditional neuron model creation paradigm (morphological reconstruction and evolutionary fit, possibly followed by a reduction), and the proposed direct experiment to reduced dendrite model paradigm. **B:** Impedance matrices and holding potentials are extracted from the response amplitude to hyper- and depolarizing current step inputs, here measured once under application of an h-channel antagonist and once under control conditions. **C:** Time-scale of largest eigenmode is extracted from average decay back to the resting membrane potential. **D:** Combined error of full model (grey) – fitted to current step data through an evolutionary algorithm, of the reduction of the full model (red) and of the direct fit of the reduced model to the data (brown). **E:** Experimentally recorded traces (grey), traces from the full reconstruction (black) and its reduction (green and purple, dashed). **F:** Experimentally recorded traces (grey) and traces from the directly fitted reduced model (green and purple, dashed).

## Discussion

We have presented a flexible yet accurate method to construct reduced neuron models from experimental data and morphological reconstructions. The method consists of linear parameter fits at holding potentials that are informative for the full non-linear dynamics. First, we derive leak and coupling conductances from the passified version of the full model. Second, we fit the maximal conductances of the active ion-channels so that the reduced model, at various holding potentials, responds similarly to input perturbations as the full model. Third, we fit the capacitances to reproduce spatio-temporal integrative properties. Finally, we ensure that resting membrane potentials in the full and reduced models match. The resulting method can be adapted to a wide variety of use cases.

A fundamental use case is deriving reduced models that retain specific elements of the dendritic computational repertoire, to explore how those dendritic computations may improve network function. To that purpose, one can apply our method to compartments placed at input sites needed for the computation of interest. For instance, if a computation requires independent computational subunits^7,41^, one would place compartments at the dendritic tips. If one wants to modulate cooperativity of excitatory inputs at distal sites, one would place compartments with shunting conductances at the bifurcations between the distal sites^9^. Detecting input sequences^10^ or implementing a dendritic integration gradient^42^ would require placing two or more compartments along the branches of interest. For many of these computations, a passive morphology with non-linear (NMDA) synapses suffices. Furthermore, our method can be combined with abstract models of AP-generation^43^.

A second use case is constructing models directly from electro-physiological recordings. Our method avoids the labour-intensive detour of reconstructing the morphology and optimizing model parameters with an evolutionary algorithm. In combination with advances in voltage-sensitive dye imaging, our method may thus form the basis of a high-throughput experiment-to-model paradigm.

A third use case is elucidating the effective complexity of a given dendritic tree. We assessed whether afferent spatial input motifs could be simplified by removing a branch or subtree and grouping the affected synapses at the next proximal compartment. We found that such simplification is possible when the input impedance of the affected synapse sites is close to the input impedance of the compartment site. If this condition does not hold, the simplification can still proceed if conductance fluctuations are small compared to the average conductance magnitude, but not otherwise. What is the simplest model for a given dendritic tree that captures its full computational repertoire? We find that spike coincidence and subthreshold voltage metrics reach satisfying accuracy at, but do not improve much beyond ~ 3 compartments per 100 μm. Further research is required to tease apart mere numerical errors from potentially missed computational features.

Computational tools have played a key role in neuroscience. Take NEURON^28^, that accelerated our understanding of morphological neurons, or NEST^44^, that enabled simulation of large-scale spiking point-neuron networks. We have implemented a Python toolbox that automatizes the simplification process (NEural Analysis Toolkit – NEAT – https://github.com/unibe-cns/NEAT). The toolbox reads any morphology in the standard “.swc” format^45^ and returns the parameters of the reduced models, while also providing tools to export the models to NEURON^28^.

Our method and toolbox fill a void in between the extremes of modelling large scale networks with abstract models, and modelling single cells in all their detail. By enabling the efficient systematic derivation of simplified neurons, amenable to simulation at the network level, our work bridges the gap between two branches of neuroscience that historically have remained separate.

## Acknowledgements

This work was supported by the European Union’s Horizon 2020 Framework Programme for Research and Innovation (Human Brain Project, Specific Grant Agreements No. 785907, SGA1-2,3) and by the SNF Sinergia grant CRSII5_180316 (with F. Helmchen). We thank all our lab colleagues for helpful discussions and critical comments on the figures.

## Author contributions

W.A.M.W. and W.S. designed the research and wrote the manuscript. W.A.M.W. performed analysis and modeling. W.A.M.W., J.J. and B.E. developed the software toolbox. U.M. and T.N. designed and performed the experiments.

## Methods

### Code availability

NEAT (NEural Analysis Toolbox), our open-source Python toolbox to obtain reduced models, is available on https://github.com/unibe-cns/NEAT.

### Morpohologies

Three exemplar cell models were used for the analysis: a cortical L2/3 Pyramidal cell^10^ (Fig 2D), a cortical L5 Pyramidal cell^12^ (Fig 2E) and a cerebellar Purkinje cell (Fig 2F). The latter morphology was retrieved from the NeuroMorpho.org repository^45^, the two others from the ModelDB repository^29^.

### Physiological parameters

In our passive models, physiological parameters were set according to Major et al^4^.: the equilibrium potential was −75 mV, the membrane conductance 100 *μ*S/cm^2^, the capacitance 0.8 *μ*F/cm^2^ and the intracellular resistance 100 **Ω** · cm. For the active models, we took ion channels and parameters according to Branco et al.^10^ for the L2/3 Pyramid, Hay et al.^12^ for the L5 pyramid and Miyasho et al.^36^ for the Purkinje cell.

AMPA and GABA synaptic input currents were implemented as the product of a conductance profile, here a double exponential^46^, with a driving force:

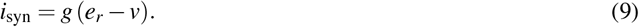

AMPA rise resp. decay times were *τ_r_* = 0.2 ms, *τ_d_* = 3 ms and AMPA reversal potential was *e* = 0 mV. For GABA, we used *τ_r_* = 0.2 ms, *τ_d_* = 10 ms and *e* = −80 mV. N-methyl-D-aspartate (NMDA) currents^47^ had the form:

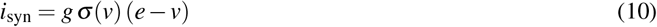

with rise resp. decay time *τ_r_* = 0.2 ms, *τ_d_* = 43 ms, and *e* = 0 mV, while ***σ***(*v*), modeling the channel’s magnesium block, had the form^48^:

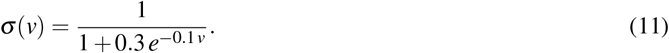

The ‘conductance’ of an AMPA or GABA synapse signifies the maximum value of its conductance window. For an AMPA+NMDA synapse, the conductance is the maximal value of the AMPA conductance window, and the conductance of the NMDA component is determined by multiplying the AMPA conductance value with an NMDA ratio *R*_NMDA_, set to be either 2 or 3.

### Biophysical models

We used the NEURON simulator^28^ to implement biophysical and reduced models. For the former, the distance step was set according the lambda rule^28^ or smaller.

### Quasi-active channels

A voltage-dependent ion channel described by the Hodgkin-Huxley formalism can in general be written as:

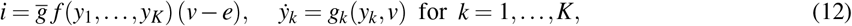

where 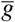 is the channel’s maximal conductance, *e* its reversal potential, *y*_1_,…,*y_K_* its state variables, *f* (.) a function that depends on the channel type (e.g. for a typical sodium channel *f* (*m, h*) = *m*^3^ *h*), and *g_k_*(*y_k_, v*) (*k* = 1,…, *K*) the functions governing state variable activation (*g_k_*(*y_k_, v*) can usually be written as (*y*_∞*k*_(*v*) – *y_k_*)/*τ_k_*(*v*) with *y*_∞*k*_(*v*) the state variable’s activation and ***τ**_k_*(*v*) its time-scale). To obtain the channel’s quasi-active approximation^49,31^ around a holding potential *v_h_* and a state variable expansion point 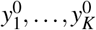, we linearise (12):

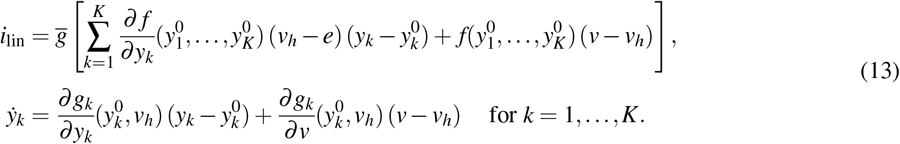

To obtain the zero-frequency contribution of (13) to the impedance and conductance matrices, we set 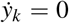 and substitute the solutions for 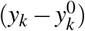 in *i*_lin_:

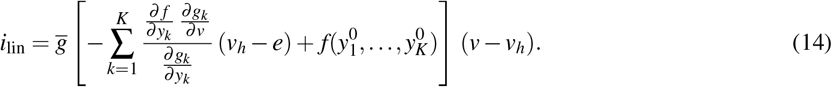

### The impedance matrix

The voltage response ***δ**v_x_*(*t*) at a location *x* along the neuron to an input current perturbation *δi*_*x*′_ (*t*) at location *x*′ can be computed as the convolution of an impedance kernel with *δi*_*x*′_ (*t*):

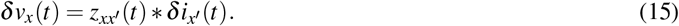

The impedance kernel itself can be computed in the frequency domain from the quasi-active cable equation using Koch’s algorithm^31^. We may assume any a-priori arbitrary set of holding potentials and ion channel state-variables to compute the quasi-active expansion (with Koch’s algorithm, the only constraint is that they be constant for each cylindrical segment). For our purpose, spatially uniform holding potentials and state-variables expansion points suffice. For a steady state current *δi*_*x*′_, (15) simplifies to:

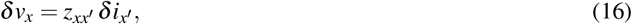

where we term 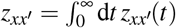 the “impedance” (*z_xx′_* is also known as the input resistance if *x* = *x′* or the transfer resistance if *x* = *x′*). For passive membranes, impedance kernels can also be computed as a sum of exponentials^33,34^:

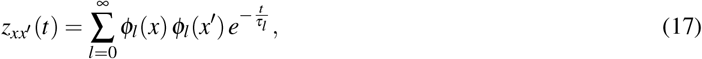

where we adopt the ordering *τ*_0_ ≥ *τ*_1_ ≥ *τ*_2_ ≥ … Essentially, this infinite sum can be thought of as the generalization of the eigenmodes for linear dynamical systems^32^ to partial differential equations.

When a current perturbation is applied at multiple sites along the neuron (we write ***δ*i**(*t*) = (***δ*i**_1_ (*t*),…, ***δi***_*n*_(*t*))), we obtain the voltage response ***δ*v**(*t*) = (***δv***_1_ (*t*),…, ***δ**v_n_*(*t*)) at those sites from:

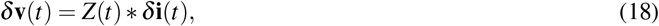

with *Z*(*t*) the matrix of impedance kernels (the *ij*’th elements of this matrix is the impedance kernel between sites *i* and *j*). In the steady state case, we call *Z* the impedance matrix.

### Compartmental models

The voltage *v_i_* in a compartment *i*, connected to a set 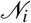 of nearest neighbour compartments, and with a set of ion channels 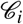 is given by

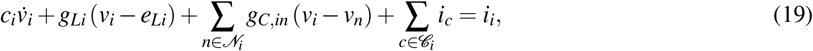

where *c_i_* is the capacitance of compartment *i*, *g_Li_* resp. *e_Li_* its leak conductance resp. reversal, *g_Cin_* its coupling conductance to the neighbouring compartment *n, i_c_* the ion channel current (12) of channel *c* in compartment *i* and *i_i_* the any arbitrary input current to compartment *i*.

### Simplification method details

For any given set of *M* sites on the morphology, we construct a simplified compartmental model whose connection structure follows a tree graph defined by the original morphology (Fig 1A). We do not allow triplet connections (e.g. sites 1-2, sites 2-3 and sites 3-1 all mutually connected) in our reduced models. Hence, to obtain accurate results it is necessary to extend the original set of M sites with the B bifurcation points that lie in between (Fig S1C). With these *N* = *M* + *B* sites, we thus define a tree graph that provides the scaffold for our fit (and our reduced model).

The fitting process proceeds in four steps: (1) fit the passive leak and coupling conductances, (2) fit the capacitances, (3) fit the maximal conductances of the ion channels and (4) fit the reversal potentials to obtain the same resting membrane potentials as in the biophysical model. The order of steps (2) and (3) is interchangeable, but not of the other steps. We describe these steps in detail below:

1. If the biophysical model contains active conductances, we compute their opening probabilities at rest and add them to the leak. Otherwise we simply take the leak as is. We then compute *Z* for the *N* compartment sites. In the passive case *Z* is independent of the holding potential. In (19) we substitute 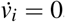. The terms *g_Li_e_Li_* are a constant contribution, and can thus be absorbed in *i_i_*. We obtain:

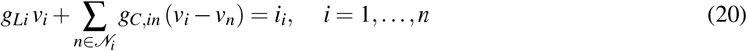

which, when recast in matrix form, yields (2). (3) then yields a system of *N*^2^ equations, linear in *g_Li_* and *g_C,in_* (note that this system always has 2*N* – 1 unknowns). We recast this system of equations in a form *A***g** = **b** and solve it in the least-squares sense for **g**.
2. To fit the capacitances, we set *i_c_* = 0 and again absorb *g_Li_ e_Li_* in *i_i_* in (19) to obtain:

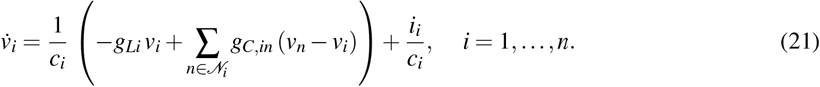 In matrix form, we obtain (5) with *S* = diag(**c**)^-1^ *G*. We require that this matrix has the same smallest eigenvalue (corresponding to the largest time-scale *τ*_0_) *α*_0_ = 1/*τ*_0_ and eigenvector *ϕ*_0_ = (*ϕ*_0_(*x*_1_),…, *ϕ*_0_(*x_N_*)) as the biophysical model, as defined by (17). Hence, we obtain the system of equations (6) that can be solved for **c**.
3. We compute the maximal conductances of each ion channel type separately. Thus, next to setting 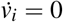, we set *i_c_* = 0 for all but one of the channels – call the non-zero channel *d* – and replace id with its quasi-active, zero-frequency approximation (14). We choose a spatially uniform holding potential *v_h_* and expansion point 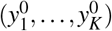 for ion channel *d*. We absorb constant terms in *i_i_* and obtain (19) in terms of *δ_v_*; = (*v_i_ – v_h_*):

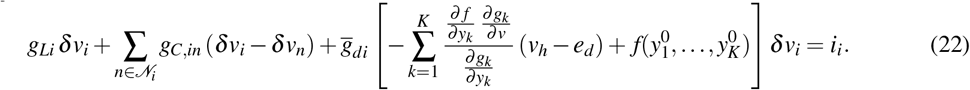 We again recast this equation in matrix form, but distinguish the known passive component *G*_pas_ from the unknown diagonal matrix containing the channel terms *G*_*v*_*h*_,chan d_:

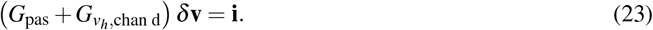 From the biophysical model, we compute *Z* by setting all ion channel conductances except channel *d* to zero, to obtain the fit equation (4). By consequence, we have a system of *N*^2^ equations, linear in the *N* unknown maximal conductances 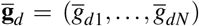, that can be recast in a form 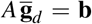, where *A* and **b** depend on the holding potential and expansion points. On the one hand, we aim to compute these linearizations for a sufficiently large range of holding potentials and expansion points. On the other hand, the fit should be restricted to domains of the ion channel phase space where the channel resides during normal input integration, so as to avoid overfitting on domains of the phase space that are never reached. We choose four holding potentials around which to linearize: *v_h_* = −75, −55, −35 and −15 mV. For channels with a single state variable *y*, we choose *y*^0^ = *y*_∞_(*v_h_*) for these four holding potentials. For channels with two state variables, we computed 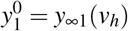 and 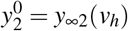 for the four holding potentials, and choose sixteen expansion points as all possible combinations of these two state variables. We then concatenate the matrices *A* and vectors **b** for each of these expansion points, and obtain the system 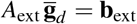 of *EN*^2^ equations (with *E* the number of expansion points) and *N* unknowns. We solve this system in the least-squares sense for 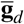 and repeat this procedure for each voltage-dependent ion channel in the biophysical model.
4. Finally we fit *e_Li_* (*i* = 1,…, *N*) to reproduce the resting membrane potential **v**_eq_ = (*v*_eq1_,…, *v*_eq*N*_) of the biophysical model evaluated at the compartment sites. We substitute this potential in (19), set 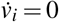 and *i* = 0 for *i* = 1,…, *N* and obtain:

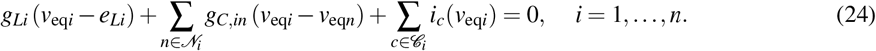 This is a system of *N* equations, linear in the *N* unknowns *e_Li_* (*i* = 1,…, *N*), and can thus be solved with standard algebraic techniques.

### Synaptic weight-rescale factors

To be able to analytically compute the weight-rescale factor when a synapse with temporal conductance *g*(*t*) is moved from its original site *s* to a compartment site *c* on a dendritic tree, we use the steady state approximation (16). Because we implicitly convolve the rescaled synaptic current with the temporal input impedance kernel *z_cc_*(*t*) at the compartment site when we simulate the reduced model, the weight-rescale factor still yields accurate temporal voltage responses (Fig 4B, E, H). In the original configuration, we have

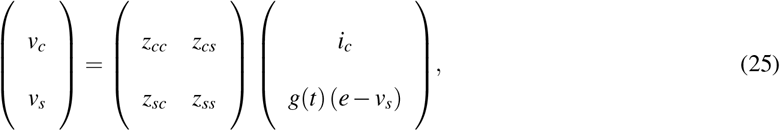

with *i_c_* an arbitrary current at the compartment site and *g* resp. *e* the synaptic conductance resp. reversal. Eliminating *v_s_* from this yields

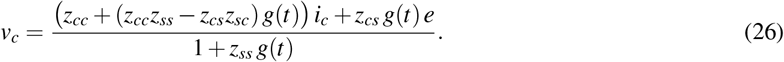

In the reduced configuration we have:

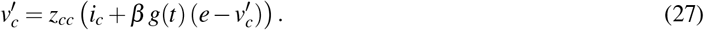

From requiring that 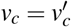, *β* is found as:

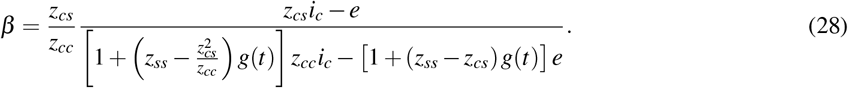

When *z_cc_ ≈ z_cs_*, which is true in basal dendrites and a reasonable approximation in many apical dendrites if *c* is on the direct path from *s* to the soma, (28) reduces to

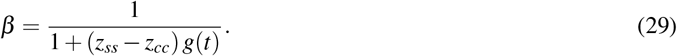

The weight-rescale factor hence depends on time. Decomposing the time-varying *g*(*t*) into *g* + *δg*(*t*), with *g* the temporal average and *δg*(*t*) the fluctuations around *g*, we find (7) and (8) from requiring that the denominator of (29) be constant in time:

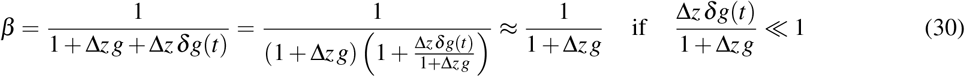

where Δ*z* = *z_ss_* – *z_cc_*.

In the case of an NMDA synapse, the conductance depends sigmoidally on the local voltage. Using scale factor (7), we obtain for the rescaled synaptic current at the compartment site:

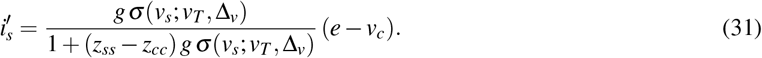

This current still depends on the voltage at the synaptic site *v_s_* in the sigmoid. A current at the compartment site that causes a voltage *v_c_* there, would have caused a voltage

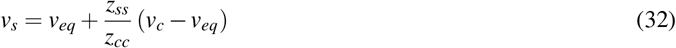

if it was placed at the synapse site. Hence, to retain the same activation level of the NMDA synapse, we substitute (32) in the sigmoid. This amounts to a change in threshold and width of the sigmoid:

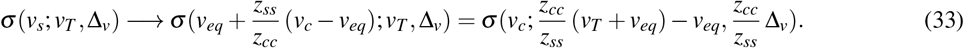

We will denote this modified sigmoid as ***σ***′ (*v*). We have not yet succeeded in reformulating (31) by only rescaling its parameters with constant rescale factors, since 1 / (1 + (*z_ss_ – z_cc_*) *g* ***σ***′ (*v_c_*)) still depends on time through its dependence on *v_c_*. To obtain constant rescale factors, we substitute the (31) in (15):

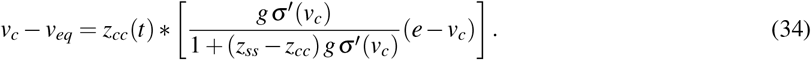

Here, the current in square brackets is the synaptic current. The convolution to obtain vc is implemented implicitly by the compartmental model. We then approximately eliminate the denominator:

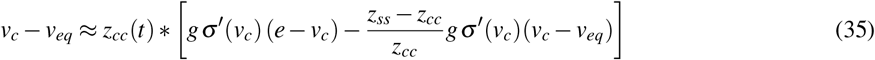

to obtain for the synaptic current:

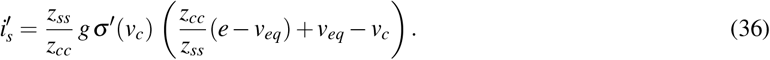

We thus find that the synaptic weight is rescaled by a constant factor 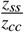 and the reversal potential shifted to 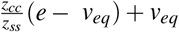. Note however that this reduction relies on two key assumptions: that there are no other inputs to the dendritic tree and that there are no fluctuations in *v_c_* generated by input currents not at site s, as this could lead to spurious activation or suppression of *σ*′(*v_c_*).

Suppose now we shift *M* conductance synapses to *N* compartments. Let **v** = (*v*_1_,…, *v_N_*, *v*_*N*+1_,…, *v*_*N+M*_) be the vector with *K* = *N* + *M* components containing the voltages at the *N* compartment sites and at the *M* synapse sites. Similarly **g** = (0,…, 0, *g*_1_,…, *g_M_*) is a vector of *K* components containing *N* zeros and the *M* synaptic conductances, while **e** = (0,…, 0, *e*_1_,…, *e_M_*) contains the synaptic reversals. In matrix form, (16) for this system becomes:

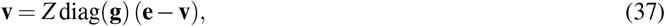

and its solution:

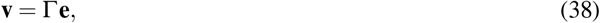

with Γ a matrix given by:

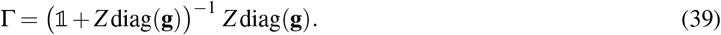

In the reduced setting, we assign each synapse to one compartment site (a reasonable choice could be the closest site). We introduce an *N × M* compartment assignment matrix *C* where an element *c_nm_* is 1 if synapse *m* is assigned to compartment *n* and zero otherwise. We further introduce a vector of reduced compartment voltages 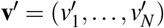 a vector of rescaled synaptic conductances 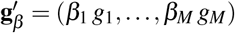 and a vector of synaptic reversals **e**′ = (*e*_1_,…, *e_M_*). In the reduced model, voltages are obtained from:

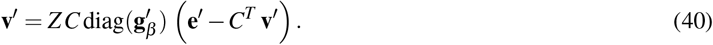

We then require that **v**_*N*_ = **v**′, with **v**_*N*_ a vector containing the first *N* components of **v**. We denote by Γ_*NM*_ the matrix containing the first *N* rows and last *M* columns of Γ, and note that we can write (38) as **v**_*N*_ = Γ_*NM*_ **e**′. Substituting this in (40) yields:

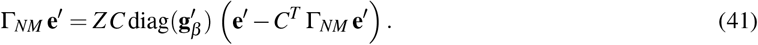

We then fit the parameters *β*_1_,…, *β_M_* (here absorbed in 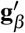) in the least-square sense from:

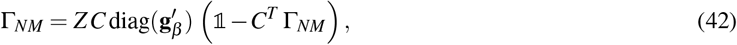

which again is a linear fit.

### Experimental recordings

Coronal slices (300 μm thick) containing the anterior cingulate cortex (ACC) were prepared from 10 – 12 week-old C57BL/6 mice using a vibratome on a block angled at 15 deg to the horizontal in ice-cold oxygenated artificial cerebral spinal fluid (ACSF) and then maintained in the same solution at 37°C for 15 – 120 min. Normal ACSF contained (in mM) NaCl, 125; NaHCO3, 25; KCl, 2.5; NaH2PO4, 1.25; MgCl2, 1; glucose, 25; CaCl2, 2; pH 7.4. Individual neurons were visualized with a Nikon Eclipse E600FN fit with a combination of oblique infrared illumination optics and epifluorescence, the switch between optical configurations was software-triggered (DanCam 2013^50^). Pyramidal neurons were selected on the clearly visible, proximal apical dendrite. This selection criterion resulted in a homogeneous population of pyramidal neurons based on their firing properties and shape of the AP (i.e., all cells possessed a prominent after-hyperpolarization and a significant sag ratio at the soma). Dual somatic and dendritic whole-cell patch-clamp recordings were performed from identified L5 pyramidal neurons in the rostroventral ACC (1.1 – 1.4 mm below the pial surface, 1.1 – 0.2 mm rostral to the Bregma) using two Dagan BVC-700. During the experiments, the external recording solution (normal ACSF) was supplemented with 0.5 mM CNQX and 0.5 mM AP-5 to block excitatory glutamatergic synaptic transmission. Experiments were performed at physiological temperatures between 34 – 37°C. Whole-cell recording pipettes (somatic, 4 to 8 MΩ; dendritic, 12 to 32 MΩ), were pulled from borosilicate glass. The internal pipette solution consisted of (in mM) potassium gluconate, 135; KCl, 7; Hepes, 10; Na2-phosphocreatine, 10; Mg-ATP, 4; GTP, 0.3; 0.2% biocytin; pH 7.2 (with KOH); 291–293 mosmol l^-1^. For somatic recordings, 10 – 20 μm Alexa 594 was added to the intracellular solution: first, the soma was patched (whole-cell configuration by negative pressure), after 5 minutes of intracellular perfusion, the fluorescent signal allowed for the clear identification of the apical dendritic tree, then the dendritic region of interest was patched with a smaller pipette. Compensation was performed in current clamp mode by recovering the fast, initial square voltage response to a hyperpolarizing current injection (−100 pA, 50 ms). First the pipette capacitance as compensated to the level that the voltage response showed an immediate voltage drop due to the series resistance of the pipette that was adjusted subsequently. Compensation of dendritic series resistance yielded values between 30 – 60 MΩ for pipettes with a resistance between 17 – 21 MΩ. On average series resistance was 2.3 times larger than the pipette resistance. Series resistance of both, somatic and dendritic recording electrodes were monitored frequently and experiments were terminated when proper compensation was not possible anymore (i.e. reached values of more than 4 times the pipette resistance). All cells were filled with biocytin and PFA-fixed slices were developed with the avidin-biotin-peroxidase method for Neurolucida reconstructions^51^. Data analysis was performed using Igor software (Wavemetrics) and Excel (Microsoft).

### Simulation-specific parameters

#### Parameters figure 2

For the sequence detection (Fig 1f), the synapse model is as in Branco et al.^10^. The maximal conductance of the AMPA component is 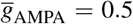 nS and its conductance window shape given by an alpha function^46^ with a time-scale of 2 ms. The NMDA component is given by a kinetic model^52^ with 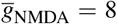 nS and external magnesium concentration of 1 mM, and receives an exponentially decaying (*τ* = .5 ms) neurotransmitter concentration with amplitude of 5 mM. For the input order detection (Fig 1G), AMPA synapses 1 and 2 use the standard parameters and have respective maximal conductances of 10 and 5 nS. For the simulation with the L5 pyramidal cell, excitatory synapses have AMPA+NMDA components with *R_NMDA_* = 2 and 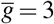 nS. For inhibitory synapses 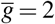 nS. In the purkinje cell, excitatory synapses only have AMPA components with 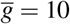 nS, while for inhibitory synapses 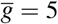 nS. For the L2/3 pyramid, L5 pyramid, resp. Purkinje cell, excitatory Poisson rates are 2, 5 resp. 6 Hz and inhibitory firing rates are 4, 1 resp. 2 Hz. Simulation were run for 10000 ms.

#### Parameters figure 3

APs are evoked with a DC current pulse. For panels A-C, pulse amplitude is *i*_amp_ = .5 nA and pulse duration is *t*_dur_ = 5 ms. For panels, D-F, we have *i*_amp_ = 3 nA and *t*_dur_ = 1 ms. For panels G-I, *i*_amp_ = 1.5 nA and *t*_dur_ = 5 ms. For panels J-K, the somatic current pulse had (*i*_amp_ = 1.9 nA and *t*_dur_ = 5 ms), while the dendritic current injection had a double exponential waveform, with *τ_r_* = .5 ms and *τ_d_* = 5 ms and amplitude *i*_amp_ = .5 nA. Onset of the somatic current pulse precedes onset of the dendritic current injection by 5 ms.

#### Parameters figure 4

In panel B, we inject OU processes for current (mean *μ* = .08 nA and standard deviation *σ* = .025 nA) and conductance (*μ* = 0.005 *μ*S, *σ* = 0.0025 *μ*S). Reversal was 0 mV for the excitatory conductance and −80 mV for the inhibitory conductance. All OU processes had a time-scale of 30 ms. In panels D-F, we use AMPA+NMDA synapses with 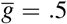 nS and *R*_NMDA_ = 3. Burst firing is mimicked by drawing synapse activation times from a Gaussian distribution with as mean the burst time and a standard-deviation of 2 ms. In panels G-J, AMPA+NMDA synapses at the compartment site (green square) have 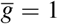 nS and *R*_nmda_ = 2. To determine the NMDA spike threshold, we activate between 0 and 200 synapses in a burst, repeat this 5 times for each number of synapses, and compute the average largest increase in voltage amplitude × waveform halfwidth. Parameters for the background AMPA and GABA synapses distributed in the apical dendrites are sampled from ranges *e_n_* ∈ [−80 mV, −50 mV], *g*_avg_ ∈ [0.01 nS,300 nS], *r*_avg_ ∈ [1Hz, 100 Hz], *n*_loc_ ∈ [1,20] and Δ*z*_avg_ ∈ [0MΩ, 1023.2 MΩ] following the latin hypercube (LH) method^53^. *g*_avg_ and *r*_avg_ are sampled on a log scale, whereas for all other parameters we use a linear scale. *e_n_* signifies the nudging potential:

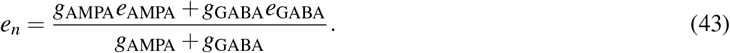

Here, *g*_AMPA_ and *g*_GABA_ are the average AMPA and GABA input conductances, found from (43) and from

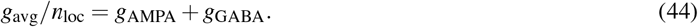

Maximal conductances of AMPA resp. GABA synapes are determined so that the time-averaged conductance, assuming a Poisson rate of *r*_AMPA_ = *r*_avg_ resp. *r*_GABA_ = 2*r*_avg_, was *g*_AMPA_ resp. *g*_GABA_.

#### Parameters figure 5

To obtain compartment sites, we divide the longest branch in each basal subtree in *n* parts of equal length, thus giving us *n* distances from the soma, where we increase *n* from 0 (point-neuron) to 10. In each subtree, we distribute compartments at all sites at these distances, and also add all bifurcation sites in between compartment sites. In this way, we obtain 11 reductions that we quantify according to ‘no. of compartments per 100 μm of dendrite.’ We implement 400 ‘background’ synapses; 200 AMPA and 200 GABA synapses with an average conductance of *g*_AMPA_ resp. *g*_GABA_ = 4*g*_AMPA_ and firing rate *r*_AMPA_ = *r*_avg_ resp. *r*_GABA_ = *r*_avg_. We implement 20 synapse clusters, consisting of AMPA+NMDA synapses with maximal conductance 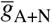 and *R*_NMDA_ = 2. These clusters are activated with a Poissonian burst rate *r*_burst_, the number of spikes per burst is drawn from a Poisson distribution with parameter *n*avg, and spike times are drawn for each burst time according to a Gaussian distribution with standard deviation of 5 ms. For each of the parameters not given previously, 10 LH samples were drawn from *g*_AMPA_ ∈ [0.4nS,0.8 nS], *r*_avg_ ∈ [1.5 Hz,3.0Hz], 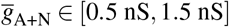, *n*_avg_ ∈ [10,30] and *r*_burst_ ∈ [0.25 Hz,0.6Hz], resulting in 10 different output spike rates shown in B. For each parameter set, a simulations was run for 10000 ms.

#### Parameters figure 6

Hyper-resp. depolarizing current steps have *i*_amp_ = −.3 nA resp. *i*_amp_ = .1 nA and *t*_dur_ = 500 ms. The full model was optimized with an evolutionary algorithm using the BluePyOpt library^14^, where we ran 100 iterations with and offspring size of 100. Goodness of fit was evaluated in a multi-objective manner as the root mean square error of the resting voltage (average voltage 100 ms before each current step), the final step voltage amplitudes after sag (average voltage during the last 100 ms of the DC current injection), and the voltage root mean square error of the full trace. We optimized the specific membrane capacitance and the conductance densities for passive leak and *h, K_ir_* and *K_m_* channels. The membrane currents followed an exponential distribution *g*(*x*) = *g*_0_*e*^*x/d*_*x*_^, with *x* the distance from the soma, and as parameters *g*_0_ – the conductance at the soma – and *d_x_* – the length constant of the distribution.

## Supplementary figures

**Figure S1.**
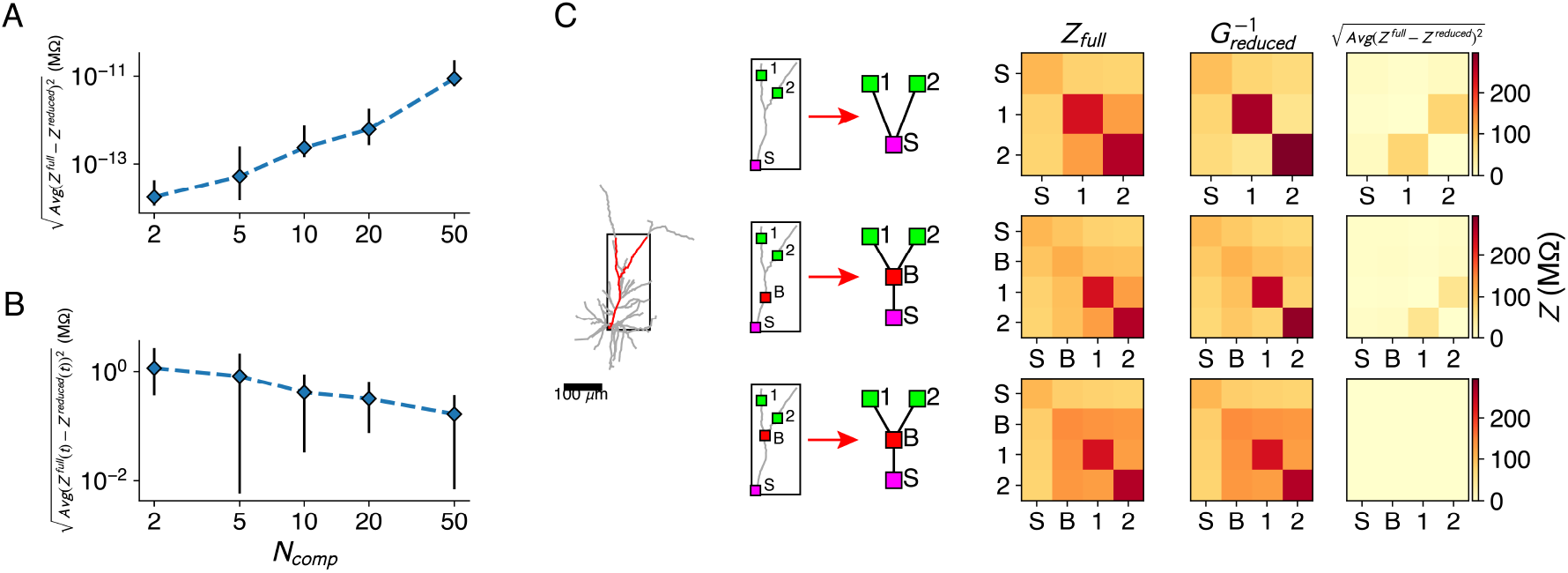
Impedance fit details. **A:** Error of the reduced model’s impedance matrix compared to the full model’s impedance matrix, for reductions of the passive L2/3 pyramid with 2, 5, 10, 20 and 50 compartments. Five random sets of fit locations were chosen for each compartment number, mean (marker) and standard deviation (error bars) are shown. **B:** Same as A, but for the impedance kernels. **C:** The bifurcations between compartment sites need to be added, as shown in the red forked dendrite (left). Soma *S* and two compartment sites 1,2 (top) were augmented with a site B on the main trunk (but not at a bifurcation, middle) or a site *B* at the bifurcation (bottom). Reductions were derived for all three configurations, but only the last configuration can accurately approximate all impedances (right).

**Figure S2.**
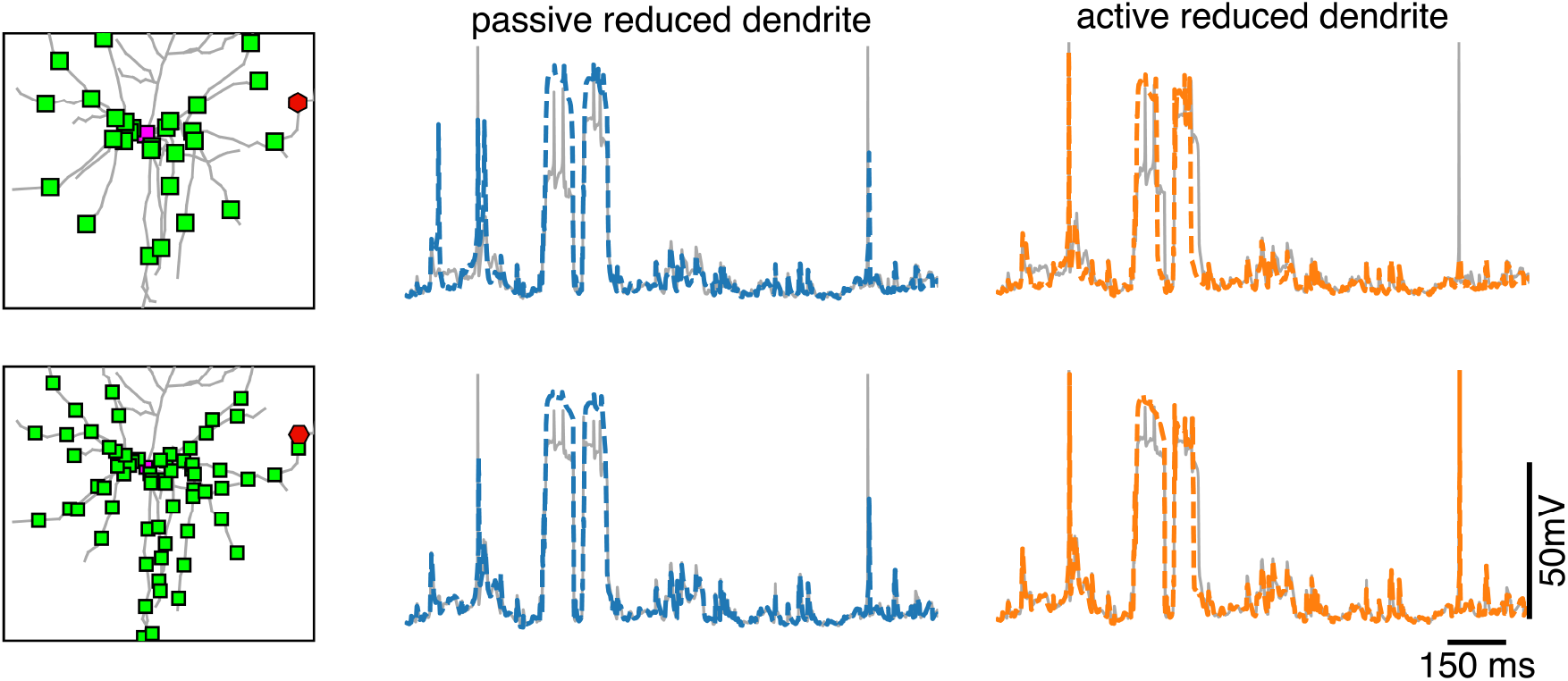
Dendritic voltage traces for reductions with passive and active dendrites. Voltage in the full model (grey) at a dendritic site (red hexagon on the left), together with the voltage in the closest compartment of the reduced model with passive dendrite (middle, blue) and of the reduced model with active dendrites (left, orange) for the same simulations as in Fig 5.

